# Seroprevalence of human Brucellosis and associated risk factors among high risk occupations in Mbeya Region of Tanzania

**DOI:** 10.1101/688705

**Authors:** Frederick D. Sagamiko, John B. Muma, Esron D. Karimuribo, Alfred A. Mwanza, Ruth L. Mfune, Calvin Sindato, Hugo Kavunga, Bernard M. Hang’ombe

## Abstract

**Background:** Brucellosis is an infectious zoonotic disease that affects humans, livestock and wildlife.

**Methods:** A cross-sectional study was conducted in Mbeya region between November 2015 and January 2016 to investigate the seroprevalence of human brucellosis and identify associated risk factors among individuals in risky occupations in Mbeya Region. A total of 425 humans from six occupational categories were serially tested for *Brucella* antibodies using the Rose Bengal Plate Test (RBPT) and competitive Enzyme Linked Immunosorbent Assay (c-ELISA), for screening and confirmation, respectively. A questionnaire survey was administered to participants collect epidemiological data.

**Results:** The overall seroprevalence among the high risk occupational individuals was 1.41% (95% CI: 0.01-0.03). Seroprevalence among the different occupations were as follows: shepherds 1.33% (95% CI: 0.14-0.22); butcher men 5.26% (95% CI: 0.10-0.17) and abattoir workers 1.08% (95% CI: 0.39-0.49). Seroprevalence was noted to vary according to occupation type, milk consumption behaviour, age and sex. Butcher men recorded the highest seroprevalence (5.0%) while individuals who consumed unboiled milk had a higher seroprevalence (1.56%) compared to those who drunk boiled milk. High seropositivity (2.25%) was observed among the age group of 1-10 years while male individuals had a higher seroprevalence (1.41%) than females (0%). Butcher men were at higher risk of exposure compared to other professions.

**Conclusion:** Our findings show the presence of brucellosis in occupationally exposed individuals in Mbeya region. There is need to sensitize the exposed individuals in order to reduce the risk of acquiring *Brucella* infections from animals and animal products This also calls for public health awareness about the disease, and implementation of control measures that will prevent further spread of brucellosis within and outside the study area..

**Author summary:** Brucellosis is a bacterial zoonosis that has evolved to establish itself as an occupational and food-borne disease Worldwide. It is responsible for huge economic losses incurred by livestock keepers and poses a public health risk to humans in most developing countries. In Tanzania, which has the 3^rd^ highest cattle population in Africa, many studies that have been done show that brucellosis exists in livestock, especially in cattle and wildlife. However, very few studies have reported on human brucellosis. The disease has been reported to occur in humans who have direct exposure to cattle or cattle products like livestock farmers, abattoir workers, veterinarians, shepherds and farm workers in many developing countries. A few studies in Tanzania have reported seroprevalences among these high-risk occupations; however, the disease has not been fully described in Mbeya region. This study was therefore aimed at filling these information gaps and contributing to the existing body of knowledge.

## Introduction

Brucellosis is a major zoonotic disease of public health and economic importance affecting domestic animals, wildlife and humans [1]. It is the second most important zoonotic disease in the world after Rabies [2]. Brucellosis is distributed worldwide but is common in countries that do not have good standardized and effective public health and domestic animal health programmes [3]. Although the genus *Brucella* consists of twelve species, it is noteworthy that this list may change as other species continue to be discovered [4]. Among the *Brucella* species, zoonotic infections are mainly attributed to *B. melitensis*, *B. abortus*, and *B. suis* [5], while *B. canis* has been mainly reported as an occupational hazard to veterinarians and laboratory workers [6]. Human brucellosis is a highly debilitating infection that presents as an acute febrile flu-like illness [7]. It is characterized by symptoms such as fever, anorexia, fatigue, headaches, depression and weight loss that may easily be confused with malaria or typhoid [7]. The source of human infection always resides in domestic or wild reservoirs.

Human cases continue to occur because of the traditional use of raw milk products and following close contact with infected animals [8, 9]. It has been observed that most cases of human brucellosis occur in rural areas where half of the people live in close proximity to their livestock, consume raw milk and make cheese using unhygienic methods [7]. Although few reports on human brucellosis exist, documentation of human cases of brucellosis in Sub-Saharan Africa is scarce, particularly reports relating to isolation of the causative agents. In sub-Saharan Africa, the prevalence of human brucellosis has been reported with varying seroprevalence ranging from 0.02% to 31.8% [10, 11, 12, 13, 14]. In Tanzania, several studies have been done in different regions including Katavi, Manyara, Morogoro, Northern Tanzania and Mwanza which have reported human brucellosis at seroprevalence ranging from 0.6 to 48.4% [15,16, 17, 18,19, 20,21]. However, there is no previous report on the disease among the high-risk human population in Mbeya region. Therefore, this study was aimed at establishing the seroprevalence and associated risk factors of human brucellosis among high-risk occupations in Mbeya region, Tanzania.

## Materials and methods

### Study area

The study was carried out in Mbeya Region in the Southern highlands of Tanzania between November 2015 and January 2016 in three selected districts namely; Mbarari, Mbeya and Momba. Geographically, Mbeya region lies about 5500 feet above sea level and experiences subtropical highland climate with humid summers and dry winters. The temperature ranges between −6°C in the highlands and 29°C on the lowlands, while the average rainfall is 900mm per year. Details of the study area have been described in our earlier publication [22]. According to the 2012 national census, the region has an estimated human population of about 2,707,[23] among which 1, 297,738 are males and 1, 409, 672 are females. A majority of the population (1, 809,298) dwell in the rural areas whereas 898, 112 are found in urban areas.

### Study population

The study population consisted of all individuals above 18 years that were involved in the cattle value chain. They were grouped into six categories; livestock professionals, shepherds, butcher men, abattoir workers, milk vendors and consumers of animal products. Sampling priority was given to individuals with direct contact/exposure to animals.

### Study design and sample size calculation

This was a cross-sectional study that was strategically designed in order to determine the seroprevalence of human brucellosis in high-risk individuals. A total of 425 humans comprising 75 shepherds from 37 *Brucella* positive herds, 11 livestock professionals, 57 butcher men, 186 abattoir workers from 4 abattoirs, 72 persons engaged in cattle milking and 24 animal product consumers were recruited in the study. The numbers of shepherds to be sampled were pre-determined from known *Brucella* positive herds based on our earlier study [22]. Among the households with infected cattle herds, only 37 were enrolled out of 53 herds. The selected study region encompassed a strategic population of individuals whose culture encourages the use of animal products for proteins, thus predisposing them to zoonotic diseases.

### Data collection

A phlebotomist aseptically collected 5ml of blood from the participant’s brachial vein using a sterile disposable syringe into pre labelled plain vacutainer tubes. The samples were then incubated overnight at room temperature and centrifuged at 3000 *xg* to get clear serum. All collected samples were assigned identification numbers and stored in a mobile refrigerator until shipment to the University of Zambia laboratory where they were stored in at −20 degree until they were examined for *Brucella* antibodies.

A pre-tested structured questionnaire was administered to the participants from whom blood was drawn in order to collect information on demographic data, socioeconomic data, exposure to animals and animal products, consumption of dairy and animal source products and the presence of specific symptoms like fever, headaches, sweats, sleeping difficulties, fatigue, weight loss, joint pain, muscle pain and back pain.

### Serological testing

#### Rose Bengal plate test

All collected sera samples were screened using Rose Bengal Plate Test (RBPT), antigen manufactured by Ubio Biotechnology Systems Pvt Ltd for detection of *Brucella* antibodies according to the test procedure recommended by OIE [1]. Briefly, 20μl of RBPT antigen and 20μl of the test serum were placed alongside on one well of the glass plate and mixed thoroughly. The slide was gently rocked for 4 minutes and thereafter, any visible agglutination was considered as a presumptive positive result.

#### Competitive Enzyme-Linked Immunosorbent Assay (C-ELISA)

RBPT positive sera were then subjected to competitive Enzyme-Linked Immunosorbent Assay (c-ELISA) as a confirmatory test, adopting a test procedure and interpretation of results as recommended by the manufacturer (Svanova Biotech AB SE-751 Uppsala, Sweden) and described by [24].

According to the ELISA kit manufacturer, serum was regarded as positive if the PI value was ≥ 30%. Only individuals that tested positive to both RBPT and c-ELISA were regarded as *Brucella* seropositive.

#### Data management and analysis

Data obtained from the serological tests and a questionnaire survey was stored in an Excel® spreadsheet database before being imported into STATA 13® statistical software for analysis. Categorical variables were summarized as frequency and percentages; continuous variables were summarized as mean or standard deviation (SD). P-values of 0.05 or less were considered statistically significant. A person was considered to be seropositive when tested positive to both RBPT and c-ELISA. The degree of association between each risk factor was assessed using the chi-square test and for all analysis, a p-value of ≤ 0.05 was taken as significant.

#### Ethical consideration

Ethical approval (reference number NIMR/HQ/R.8a/Vol.1X/2050) was obtained from the Medical Research Committee of the United Republic of Tanzania prior to the study. Individual written consent was obtained from guardians for individuals that were less than 18 years prior to enrolment. Informed consent was obtained from all participants using written and verbal explanation of the study purpose and procedure using the Swahili language.

## Results

A total of 425 individuals working in the cattle value chain in Tanzania were included in the study (Table 1). The overall human brucellosis prevalence was 1.41% (95% CI: 1.7-2.6). No female participant (n=334) tested positive for brucellosis even though these were in the majority compared to males (n=91). *Brucella* seroprevalence was recorded in three occupational categories out of the six that were considered in our study (Table 2). It was also observed that 75.3% of respondents (n=320) consumed raw milk and only 24.7% (n=10) consumed boiled milk and none had a history of consuming pasteurized milk (Table 2). The predictor variables were assessed for collinearity using Pearson’s Chi-square test and revealed a strong association between the occupation of an individual and his/her sex (P-value=<.0001) and age category (P-value=<.0001).

**Table 1:**
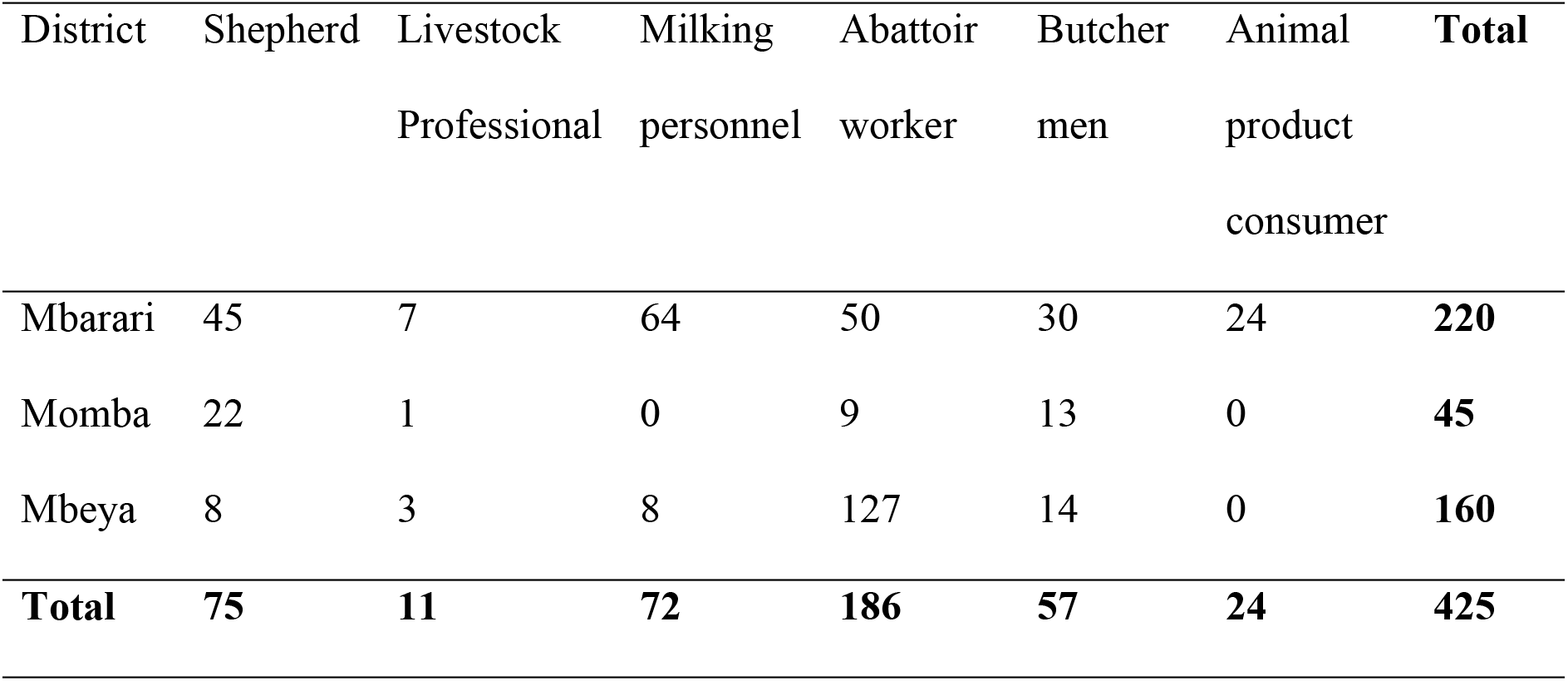
Human blood sample distribution by study district and occupation in Mbeya Region.

**Table 2:** Results of univariate analysis of seroprevalence of human brucellosis by different variables.

| Variable | Level | No | Distribution | Prop.<br>positive<br>(%) | 95% CI | P-value |
| --- | --- | --- | --- | --- | --- | --- |
| Sex | Male | 91 | 21.4 | 1.41 | 0.18-0.26 | 0.19 |
|  | Female | 334 | 78.6 | 0.00 | 0.74-0.82 |  |
| Area | Mbarari | 220 | 51.8 | 1.36 | 0.47-0.57 | 0.64 |
|  | Momba | 45 | 10.6 | 0.00 | 0.08-0.14 |  |
|  | Mbeya | 160 | 37.6 | 1.88 | 0.33-0.42 |  |
| Age | 1-10 yrs | 159 | 37.4 | 2.25 | 0.33-0.42 | 0.27 |
|  | 11-20 yrs | 196 | 46.1 | 1.02 | 0.41-0.51 |  |
|  | 21-30 yrs | 65 | 15.3 | 0.00 | 0.12-0.20 |  |
|  | 31-40 yrs | 5 | 0.1 | 0.00 | 0.00-0.03 |  |
| Occupation | Shepherd | 75 | 17.6 | 1.33 | 0.14-0.22 | 0.17 |
|  | Livestock officer | 11 | 2.5 | 0.00 | 0.01-0.05 |  |
|  | Butcher men | 57 | 13.4 | 5.26 | 0.10-0.17 |  |
|  | Abattoir worker | 186 | 43.7 | 1.08 | 0.39-0.49 |  |
|  | Milking | 72 | 17.0 | 0.00 | 0.14-0.21 |  |
|  | Animal product | 24 | 5.6 | 0.00 | 0.04-0.08 |  |
| Source of milk | Cattle | 425 | 1 | 1.41 | - | - |
|  | Goat | 0 | - | 0.00 |  |  |
| Milk consumption | Unboiled | 320 | 75.3 | 1.56 | 0.71-0.79 | 0.65 |
|  | Boiled | 105 | 24.7 | 0.95 | 0.21-0.29 |  |
| Consuming raw blood | No | 425 | 1 | 1.41 | - | - |
|  | Yes | 0 |  |  |  |  |
| Assisting parturition | No | 312 | 73.4 | 1.60 | 0.69-0.77 | 0.58 |
|  | Yes | 113 | 26.6 | 0.88 | 0.23-0.31 |  |

### Seroprevalence of human brucellosis

There was no statistical association between *Brucella* seropositivity and all the hypothesized risk factors evaluated in univariate analysis (Table 2). However, results indicated an apparent variation of seroprevalence by occupation, milk consumption behavior, age and sex. Butcher men recorded the highest seroprevalence (5.0%) while a higher seroprevalence (1.56%) was recorded among individuals who consumed unboiled milk compared to drinkers of boiled milk (Table 2). High seropositivity (2.25%) was observed among the
 age group of 1-10 years while male individuals had a higher seroprevalence (1.41%) than females (0%) as shown in Table 02.

## Discussion

Brucellosis is among the diseases categorized as a neglected zoonosis by the WHO. This is so because, despite its wide distribution and potentially harmful effect on human health, it has not been given due attention compared to other diseases. Generally, the public health importance of brucellosis is acknowledged throughout the world; however people in certain occupations or settings still face increased risk of exposure to the *Brucella* pathogen. These may include many players in the livestock value chain starting from livestock keepers, veterinarians/livestock officers, animal handlers, laboratory workers, slaughterhouse workers, butcher-men and consumers of animal products (meat, milk, geese).

The aim of this study was to estimate the seroprevalence of human brucellosis and identify associated risk factors among individuals engaged in risky professions in Mbeya Region. The study found an overall seroprevalence of 1.41% among shepherds, butcher men and abattoir workers in Mbeya region. This seropositivity is lower than the 2.2% reported [1] in the Kilimanjaro region and the 48.4% reported by [20] in Mwanza region. The difference can be attributed to differences in geographical locations and the use of a single but highly sensitive test (Rapid agglutination test) in the previous study. In other parts of Africa, brucellosis studies among high risk occupations in Ghana and Nigeria in slaughterhouses found seroprevalences of 9.6% [25] and 24.1% [11] while a study in Sudan found varying prevalences of 9.5%, 15.3%, 24.4% and 26.5% among veterinarians, meat inspectors, abattoir workers and animal handlers, respectively [26]. The findings in Ghana, Nigeria and Sudan were higher than those found in this study in all occupational groups. These result does not justify the lower levels of human brucellosis in our study area as most of the people had a tendency of taking antibiotics regularly [27]. In so doing, they could treat brucellosis unknowingly and thereby negatively cause the existing problem of antimicrobial drug resistance. Brucellosis has also been reported in slaughterhouse workers in Iran with the prevalence of 7.8% [28] and 9.8% [9]. In Lahore district of Pakistan, the prevalence of brucellosis in abattoir workers was found to be 21.7%, which was higher compared to that observed in our study area [30]. This can be explained by the fact that both Pakistan and Iran depend on sheep and goat meat and milk for protein sources, which are likely to be contaminated with *B. melitensis*. In humans, *B. melitensis* is highly infectious compared to *B. abortus* [31], which is likely to be the problem in the study area. This practice followed in marketing and distribution of sheep and goat meat as well as milk products, in particular, makes the enforcement of hygienic measures very difficult [7].

### Risk factors associated with *Brucella* seropositivity

The prevalence of human brucellosis in occupationally exposed individuals in the Mbeya region of Tanzania has been noted to vary with the occupational category, milk consumption behaviour, age and sex, although this was not statistically significant. The butcher men had a higher risk of exposure to brucellosis than shepherds and abattoir workers. This could be attributed to that fact that there is increased risk of injury (knife-cuts) among the butcher men compared to the other categories, thus increasing the exposure risk to the Brucella pathogen. High prevalence of brucellosis in males can be explained by the fact that most of activities in cattle value chain are carried out by males than females. Since majority of butchermen are males who has high risk of acquiring brucellosis, it can be the reasons for the high prevalence of the disease. This is similar to findings by other studies [32, 33]. However, none of these have establish the risk that shepherds have towards brucellosis in Tanzania despite the fact that they are at the starting point of the livestock value chain. Therefore, this is the first report in Tanzania to establish the risk that shepherds have towards brucellosis. These findings are in contrast to those by [20] who reported a high risk of exposure to brucellosis abattoir workers. The high disease prevalence among butcher men could be because they spend longer periods handling animal carcasses usually without protective wear and are more likely to be injured when cutting meat and are in close contact with blood and tissues of infected animals. Hence they are at higher risk of infection than other groups. *Brucella* seropositivity was higher in males (1.41%) than females (0%), similar to findings by [11] in Nigeria but contrary to findings by [17] in Morogoro. Seroprevalence was higher in individuals below 20 years of age (2.25%). These findings agree with those by [34] in Uganda. This can be attributed to the traditional role that young men play in livestock management among the pastoral communities. The young males start rearing livestock at a young age and are in direct contact with animal and animal products during their daily livestock activities, which increases their risk of exposure to brucellosis. In our study, 75.3% of people consumed raw milk which is higher than that reported by [35]. This can be explained by the fact that over 70% of milk sales in Tanzania is produced by pastoral farmers who do not believe or know that milk could be a potential source of infection to humans; and are not ready to subject their milk to any form of [36]. Milk is a major vehicle for transmission of *Brucella* infection and individuals with a history of consumption of raw milk were more likely to be infected [34].

## Conclusion and Recommendations

Our findings demonstrate the presence of human brucellosis in occupationally exposed individuals, specifically abattoir workers, butcher men and shepherds in the Mbeya region. Given that we applied random sampling strategy to obtain the sample size (from our earlier study), the study findings can be generalized to the region. The results from this study indicate that more work needs to be done to educate the occupationally exposed individuals on brucellosis and its associated risks. Therefore, there is need to create public awareness, design and implement control measures that will prevent further spread of the disease within and outside the study area. We recommend regional and multi-sectoral collaboration, especially among veterinarians and medical professionals using the one health approach in order to combat the disease.

## Limitations encountered in this study

Some of the limitations were that some shepherds from certain cattle herds that had been screened earlier could not be screened due to the migratory nature of agro-pastoralism in search of water and pasture.

## Acknowledgement

The authors are grateful to Mr Joseph Ndebe, Ms Jessica Chitambo for assisting in laboratory work, Health Department of Mbarara, Momba and Mbeya District Councils for their assistance during the sampling.

## Author contributions

**Conceptualization:** Frederick D. Sagamiko and John B. Muma

**Data curation:** Frederick D. Sagamiko

**Formal analysis:** Frederick D. Sagamiko, John B. Muma and Bernard M. Hang’ombe

**Funding acquisition:** INTRA-ACP Mobility Project

**Methodology:** Frederick D. Sagamiko

**Supervision:** John B. Muma, Bernard M. Hang’ombe, Alfred M. Mwanza.

**Validation:** Bernard M. Hang’ombe

**Visualization:** Frederick D. Sagamiko

**Writing – original draft:** Sagamiko FD, John B. Muma, Bernard M. Hang’ombe, Esron D. Karimuribo, Alfred M. Mwanza, Ruth L. Mfune, Calvin Sindato, Hugo Kavunga.

**Writing – review & editing:** John B. Muma, Bernard M. Hang’ombe, Esron D. Karimuribo, Alfred M. Mwanza, Ruth L. Mfune, Calvin Sindato, Hugo Kavunga.

## Supporting Information

**S1 Checklist. STROBE Checklist** (DOC)

**S2 Ethical Approval. Approval letter from Ministry of Health (MOH), Tanzania for collection of human blood samples** (PDF)

**S3 Dataset. Human dataset used for analysis** (XLS)

